# Interpretable deep learning framework for understanding molecular changes in human brains with Alzheimer’s disease: implications for microglia activation and sex differences

**DOI:** 10.1101/2023.12.18.572226

**Authors:** Maitry Ronakbhai Trivedi, Amogh Manoj Joshi, Jay Shah, Benjamin P Readhead, Melissa A Wilson, Yi Su, Eric M Reiman, Teresa Wu, Qi Wang

## Abstract

**INTRODUCTION:** The objective of this study is to characterize the dysregulation of gene expression in AD affected brain tissues through an interpretable deep learning framework.

**METHODS:** We trained multi-layer perceptron models for the classification of neuropathologically confirmed AD vs. controls using transcriptomic data from three brain regions of ROSMAP study. The disease spectrum was then modeled as a progressive trajectory. SHAP value was derived to explain model predictions and identify significantly implicated genes for subsequent gene co-expression network analysis.

**RESULTS:** The models achieved excellent performance in classification and prediction in two external datasets from Mayo RNA-seq cohort and Mount Sinai Brain Bank cohort. SHAP explainer revealed common and specific transcriptomic signatures from different brain regions.

**DISCUSSION:** We identified common gene signatures among different brain regions in microglia and sex specific modules in neurons implicated in AD. This work paves the way for utilizing artificial intelligence approaches in studying AD at the molecular level.

**Research-in-Context:**

1. Systematic review: Postmortem brain transcriptomes have been analyzed to study the molecular changes associated with Alzheimer’s disease, usually by a direct contrast approach such as differential gene expression analysis. Nuanced gene regulatory networks thus cannot be easily pinpointed from convoluted data such as those from bulk-tissue profiling. We applied a novel interpretable deep learning approach to dissect the RNA-seq data collected from three different brain regions of a large clinical cohort and identified significant genes for network analysis implicated for AD.
2. Interpretation: Our models successfully predicted neuropathological and clinical traits in both internal and external validations. We corroborated known microglial biology in addition to revealing novel sex chromosome-linked gene contributing to sex dimorphism in AD.
3. Future directions: The framework could have broad utility for interpreting multi-omic data such as those from single-cell profiling, to advance our understanding of molecular mechanisms of complex human disease such as AD.

**Highlights:**

- We applied novel interpretable deep learning methods to postmortem brain transcriptomes from three different brain regions
- We interpreted the models to identify genes most strongly implicated in AD
- Network analysis corroborated known microglial biology and revealed novel sex specific transcriptional factors associated with neuronal loss in AD

## 1. Background

Despite extensive research of Alzheimer’s disease (AD) for its underlying mechanism of onset, manifestation, and progression, the complex molecular events behind the disease spectrum remain incompletely understood[1]. Recently, large-scale high-throughput profiling of omics data such as RNA-seq has enabled the application of novel machine learning (ML) methods to dissect the gene expression profiles of postmortem brain tissues from large clinical AD cohorts, thus opening the channel of artificial intelligence (AI) approach for advancing our understanding and seeking potential early treatment of the devastating disease[2].

One of the challenges faced by the application of AI approaches to multi-omic data is lack of interpretability. Complex ML models, like Deep Neural Networks (DNNs), although with unparalleled predictive power, are often considered as “black box” models as their decision-making processes are not easily inspected by human investigators[3]. Existing literature has reported the use of different AI frameworks to uncover deep interrelationships between gene expression and AD neuropathologies[4, 5]. However, due to the limited sample sizes available in these studies, the interpretation of the models, specifically the DNNs, had to be oversimplified, and relied on outcome trained from datasets aggregated across multiple brain regions, which are known to be affected by AD neuropathology quite differently[6]. Achieving the full potential of modern AI techniques in genomics research necessitates a thorough evaluation of the model, and a precise, in-depth, and comprehensive interpretation of its mechanisms based on input features. Most recently, through the Accelerating Medicines Project for Alzheimer’s Disease (AMP-AD) Target Discovery Consortium and associated open-science consortia[7], multidimensional molecular data from more than 2,000 human brains and peripheral tissues from multiple AD cohorts have been made publicly available [8]. Coupled with comprehensive neuropathological and clinical characterizations, these data represent an unprecedented opportunity to advance our understanding of the molecular basis of AD.

Herein we report efforts to build and interpret DNN models based on our published ML framework[4] using bulk RNA-seq data from three different brain regions in the Religious Orders Study and Memory and Aging Project (ROSMAP) cohort[9, 10] from the AMP-AD open data platform, as illustrated in Figure 1. We extend our previous work by applying the framework to data from multiple brain regions with larger sample sizes, and utilizing state-of-the-art model interpretation methods[11] to obtain novel biological insights. The models achieved superior performance in correlating expression profiles with neuropathological and clinical traits, and demonstrated satisfying prediction accuracy when applied to two independent AMP-AD datasets, the Mayo RNA-seq study cohort (MAYO)[12] and the Mount Sinai Brain Bank (MSBB) study cohort[13]. We identified gene modules associated with microglia activation within multiple brain regions and a region and sex specific transcription factor linked with sexual dimorphism in AD. We believe that this work lays the foundation for the application of explainable artificial intelligence approaches to high-dimensional molecular data and thus advance the study of AD etiology.

**Figure 1.**
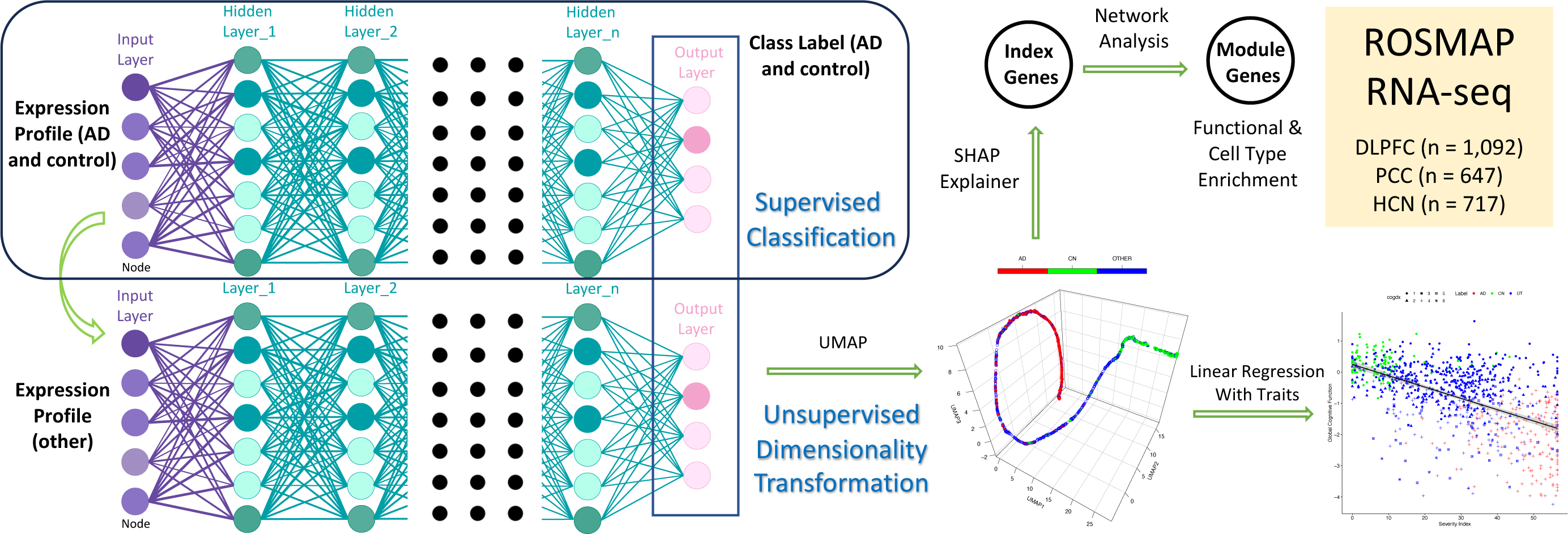
The deep learning and interpretation framework employed in this work. 1) Model training. Using the gene expression profiles from AD and control subjects and their diagnosis class as the input for supervised classification, the model was trained by a multilayer neural network. The trained network was passed forward to the profiles from the whole cohort with the resulting output manifold subject to unsupervised dimensionality transformation (UMAP) to obtain the pseudo-temporal trajectory and SI. SI was linearly correlated with phenotypic data for evaluation. 2) Trained model was interpreted by SHAP explainer to obtain the most salient features (IGs). Their co-expression relationship was examined for biological interpretation. The framework was applied to the three brain regions (DLPFC, PCC, and HCN) from ROSMAP cohort respectively.

## 2. Materials and Methods

### 2.1 RNA-seq datasets from AMP-AD consortium

All RNA-seq data were obtained from the AMP-AD data portal through Synapse (https://www.synapse.org/). Demographic information for each of the cohort (ROSMAP, MAYO and MSBB) sampled in the RNA-seq study is reported in Supplementary Table 1. The processed, normalized data were obtained for each cohort respectively, from the harmonized, uniformly processed RNA-seq data set across the three largest AMP-AD contributed studies (syn21241740). In the ROSMAP studies (syn3219045 and syn22695346), the brain tissue samples were collected from three different brain regions: dorsolateral prefrontal cortex (DLPFC, n = 1,092), posterior cingulate cortex (PCC, n = 647), and head of caudate nucleus (HCN, n = 717). In Mayo RNA-seq study (syn5550404), brain tissue samples were collected from cerebellum (CER, n = 246) and temporal cortex (TCX, n = 259). The MSBB study (syn3159438) has over 1,000 samples from the Mount Sinai/JJ Peters VA Medical Center Brain Bank, which were sequenced from 312 subjects from four brain regions including the frontal pole (FP, Brodmann area (BM) 10, n = 304), inferior frontal gyrus (IFG, BM 44, n = 297), superior temporal gyrus (STG, BM 22, n = 324) and parahippocampal gyrus (PHG, BM 36, n = 308), respectively. The harmonized processing of all the data from the three cohorts was accomplished using a common workflow (https://sage-bionetworks.github.io/sageseqr/).

The conditional quantile normalized[14] log counts per million reads (CPM) values from each data set (syn26967453, syn27024965, and syn27068756) were used in all the subsequent analyses.

### 2.2 Phenotypical data

The detailed definitions of phenotypic measurements used in the study, including clinical evaluation and postmortem neuropathological quantifications, together with their possible values of all three cohorts are reported in Supplementary Table 2.

All the clinical and pathological data for the ROSMAP cohort were obtained from the Rush Alzheimer’s Disease Center Research Resource Sharing Hub (https://www.radc.rush.edu/), upon approval of data-usage agreement. The details of the variables can be found in Supplementary Table 2.

For MAYO and MSBB cohorts, subject clinical and pathological data were obtained from Synapse (syn27000373 and syn23277389 for Mayo samples and syn27000243 for all the MSBB samples). For MAYO cohort, the following data were used in the linear regression to validate the two phenotypes: Target variables: Braak = Braak stage; Thal = Thal amyloid stage. Dependent variables: ageDeath = age at death; sex = sex; race = racial group; apoe4 = apoe4 allele count; RIN = RNA integrity number; PMI = post-mortem interval. For MSBB cohort, the following data were used in the linear regression to validate the four phenotypes: Target variables: Braak = Braak stage; plaqueMean = plaque mean density; CDR = clinical dementia rating; CERAD = CERAD (Consortium to Establish a Registry for Alzheimer’s Disease) score. Dependent variables: ageDeath = age at death; sex = sex; race = racial group; apoe4 = apoe4 allele count; RIN = RNA integrity number; PMI = post-mortem interval. The original CERAD score in the MSBB cohort was defined as: 1, Normal; 2, Definite Alzheimer’s disease; 3, Probable Alzheimer’s disease; 4, Possible Alzheimer’s disease. They were recoded to be semi-quantitative as follows: 1, Definite Alzheimer’s disease; 2, Probable Alzheimer’s disease; 3, Possible Alzheimer’s disease; and 4, Normal, to be consistent with the notion used in the ROSMAP cohort. APOE genotypes were obtained from whole genome sequencing harmonization (syn11707420) whenever it is missing in the original meta data file. Semiquantitative measurements (e.g. Braak stage, Thal phase, CERAD score, etc) were treated as quantitative. Quantitative measurements (e.g. amyloid or tangles) were log transformed.

### 2.3 Deep learning of the transcriptome from three brain regions in ROSMAP cohort

We applied a modified ML framework from our previous publication[4] to the datasets in ROSMAP cohort for the three brain regions respectively. The framework consists of two major components, supervised classification (deep learning) and unsupervised dimensionality transformation (Figure 1).

For the DNN classification, we trained multi-layer perceptron (MLP) models using neuropathologically confirmed AD patients and normal controls (CN), the two termini of the AD spectrum to maximally differentiate the two groups. Data collected from ROSMAP including cogdx, braaksc and ceradsc were used to define the class label for AD, control (CN) and OTHER groups:

1. AD: cogdx = 4, braaksc ≥ 4 and ceradsc ≤ 2;
2. CN: cogdx = 1, braaksc ≤ 3 and ceradsc ≥ 3;
3. OTHER: all the other samples.

Total number of samples used in the training for each model, their diagnosis classifications and split by training/test sets for a five-fold cross validation, and the model configurations are reported in Supplementary Table 3. For the DLPFC dataset, visualization of gene expression of sex chromosomes (XIST as X-chromosome marker and UTY as Y-chromosome marker) revealed two samples with high expressions of both markers, and they were subsequently removed from the modeling process. The input logCPM values from AD and CN samples were feature-wise z-transformed, and the mean and standard deviation (SD) of each feature were used for scaling for other samples/datasets used in validations.

For all the model training, we adopted the Adam optimizer[15], with a learning rate set to 0.0001, to update the models’ parameters. The training consisted of 300 epochs and for the purpose of selecting the most suitable model, a model checkpoint callback was used to store the weights of the best model based on the validation accuracy. The performance evaluation of these models was carried out using a comprehensive set of classification metrics, including test accuracy, sensitivity/recall, specificity, F1 score, area under the receiver operating characteristic curve (AUC) and precision[16].

After each MLP model was trained, the data samples labeled as OTHER were scaled using the mean and SD values from the AD-CN samples used in the training process and added back to the dataset. Eventually a manifold representation was generated through a forward pass utilizing the previously trained neural networks to all the samples of the whole cohort. To visualize the manifold, the dimensions of resulting representation of the final layer was further reduced to a three-dimensional UMAP space, by the “umap.UMAP.fit_transform” function in python, with the following parameters: n_components = 3, metric = “Euclidean”. Other parameters are tuned to be specific for each brain region as reported in Supplementary Table 3.

### 2.4 Applying the deep learning model to external data sets (MAYO/MSBB)

The harmonized, uniformly processed RNA-seq data sets from MAYO and MSBB were first sorted by the same gene order as the input data set of ROSMAP. Batch effects were then removed by the ComBat[17] function in the R package sva[18]. The input expression matrix subsequently was transformed to Z-score by scaling to the training dataset in the ROSMAP deep learning model. A manifold representation was obtained for all the samples in each cohort by forward pass of the trained network. Trajectories were obtained by carrying out the UMAP transformation of the existing embedding model from each set of ROSMAP data, by the “umap.UMAP.fit_transform” function using the same parameters as each model in python.

### 2.5 Model validation by correlation with phenotypic data

We derived an index, the Severity Index (SI), for staging the progression of AD from normal control to terminal disease based on the pseudotemporal trajectory in the UMAP embedding. SI was derived for each sample by applying the method of inferring pseudotimes for single-cell transcriptomics from the function ‘slingPseudotime’ as implemented in the R package Slingshot[19]. To evaluate the models built from three brain regions respectively, SIs were then linearly correlated with all the AD clinical and pathological biomarkers individually, in both the ROSMAP cohort and the two other independent cohorts (MAYO/MSBB), including the covariates PMI, RIN, apoe4 allele count, age at death, sex, race, and educ (ROSMAP only), using the following linear regression model:

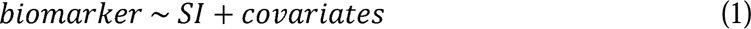

For ROSMAP, we also included the following non-AD neuropathological measurements as additional covariates: r_pd, r_stroke, dlbdx, hspath_typ, arteriol_scler, tdp_st4, caa_4gp, cvda_4gp2, ci_num2_gct, and ci_num2_mct. Their detailed definitions can be found in Supplementary Table 2 and data collection is reported in Tasaki et al[20]. The target dependent variables (biomarkers) are the AD neuropathological and clinical measurements, also reported in Supplementary Table 2. In MSBB cohort, duplicated samples were sequenced for the same individuals in some regions, and only one sample with the lowest rRNA rate was kept in the linear regression model. Correlation coefficients were obtained by the ‘lm’ function in R. The proportion of variance explained (PVE) for each predictor was obtained from the incremental sums of squares table by the ‘anova’ function in R on the model, using the order as reported.

### 2.6 Model interpretation

We applied the SHAP (SHapley Additive exPlanations)[21] tool to interpret the MLP model trained for each brain region, for extracting gene features that explain the classification. The following steps were performed to derive a quantitative SHAP metric for each feature:

1. The SHAP values for each feature *x_i_* were computed using the “shap.Explainer” function from the SHAP library in Python.
2. For each feature in each sample, calculate the SHAP values for two diagnosis classes separately:
  a. Two sets of SHAP values (*SHAP_AD_* (*x_i_*) and *SHAP_CN_*(*x_i_*)) were derived separately for AD and CN samples, where i ∈ [1,…,N_features_], N_features_ represents the total number of features.
  b. *SHAP_AD_* (*x_i_*) represents the SHAP values for all AD samples for the *i^th^* feature
  c. *SHAP_CN_*(*x_i_*) represents the SHAP values for all CN samples for the *i^th^* feature
3. For each class, compute the mean SHAP values for each feature. We computed the mean SHAP values for each feature by averaging the individual SHAP values across the samples in the AD and CN classes separately:
  a. For each feature *x_i_*, computed the mean SHAP value for AD samples as equation (2), where *N_AD_* represents the total number of AD samples in the dataset.

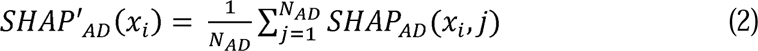
  b. For each feature *x_i_*, computed the mean SHAP value for CN samples as equation (3), where *N_CN_* represents the total number of CN samples in the dataset.

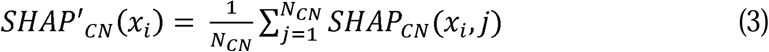
4. Compute the mean absolute SHAP Value of each feature for the model:

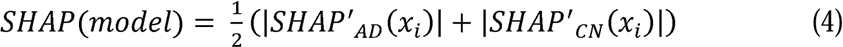
5. Run an ensemble of 100 models, each with a different random seed and the same hyperparameters, with the goal of sampling a wide range of feature relevance across different model iterations. The mean SHAP values of all 100 models were then evaluated as the final SHAP value for each feature:

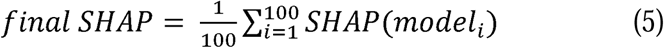

Eventually, we sorted the features by their final SHAP values calculated in equation (5) in descending order to identify the most influential features for the classification model while making predictions. The cutoff to extract features making significant contributions to the classification was set as 1.5 * interquartile range (IQR) above the third quartile of the distribution. These gene features were eventually identified as the “index genes” in each brain region that explains AD vs control classification.

### 2.7 Gene network and functional analysis

Gene co-expression networks were built by MEGENA[22], based on the expression values of the “index genes” from the residualized counts data after regressing out significant covariates, with three covariates (diagnosis, age and sex) added back, for the three brain regions (syn31141704) respectively. Common networks between those modules from different brain regions were identified by CoDiNA[23], using the correlation coefficients of significant correlation pairs derived by MEGENA in each region. Common networks in another brain region were deemed to overlap with a DLPFC submodule by Fisher’s Exact test, using the union of the IGs from the two regions as the background.

Functional and cell type enrichment analysis for each module identified in DLPFC networks was performed using Metascape[24], which uses a hypergeometric test and Benjamini-Hochberg P-value correction to identify ontology terms that contain a statistically greater number of genes in common with an input list than expected by chance, using the whole transcriptome as background. Statistically significant enriched terms based on Gene Ontology[25], KEGG[26], Reactome[27] and MSigDB[28] were clustered based on Kappa-statistical similarities among their gene memberships. A 0.3 kappa score was applied as a threshold to identify enriched terms.

All the gene enrichment analyses were performed in R (version 4.0.0)[29] by Fisher’s Exact Test (FET) on the overlaps between the gene sets of interest, using the whole transcriptome as background.

Transcriptional gene regulators were identified by Trena[30], using the expression profile of the neuron co-expression module in DLFPC in combination with those transcription factors with known motifs[31, 32]. Reprocessing of all the ROSMAP data to include combined ZFX+ZFY gene counts was accomplished following the same workflow as previous reported (https://sage-bionetworks.github.io/sageseqr/), excluding four samples with ambiguous sex markers expression. The conditional quantile normalized log counts per million reads (CPM) values were used in the analyses.

## 3. Results

### 3.1 Classification model architecture and performance

We initiated our ML framework with a DNN classification model of the two termini of the AD disease spectrum, with the goal of separating the AD patients from normal control participants as much as possible. Samples were partitioned into training and testing sets, containing 80% and 20% of the samples respectively. The detailed information about the number of AD and CN samples for each brain region is shown in Supplementary Table 3. Based on our experiments and literature reports[33], the final model utilized for this classification task in each brain region was an MLP with their respective architecture reported in Supplementary Table 3. After fine tuning of all hyperparameters, final model performance was evaluated by all classification metrics (reported in Supplementary Table 3), with receiver operating characteristic (ROC) curves shown in Supplementary Figure 1. Notably, the model trained from DLPFC (DLPFC model for simplicity, the same below) exhibited exceptional classification accuracy, achieving a test accuracy of 97.8% and a sensitivity of 100%, indicating its superior proficiency in accurately identifying AD samples. Conversely, the PCC model exhibited a comparably high testing accuracy of 96.0% and a sensitivity of 96.2%, highlighting its strong classification capabilities. The HCN model performed the least satisfactorily in comparison to the cortex dataset (DLPFC and PCC) models, showing a testing accuracy of 81.1% and a lower specificity of 56.3%. The distinction of the performance between these models also aligns with the variation in relevance to AD pathology among these brain regions, as the cortex regions are known to be highly associated with AD in contrast to subcortical regions[34].

### 3.2 Model the whole AD spectrum as a pseudotemporal trajectory

Given that AD is a progressive disease, we reasoned that the application of a pseudotemporal trajectory approach might yield insight. After classifying the two termini of the disease spectrum from their expression profiles, we projected all samples from the whole cohort to the same UMAP space by passing expression data into the trained model. As expected, we observed that the samples from AD/CN are mainly located at each of the two termini of a trajectory as two distinct clusters (Fig. 2A, C, and E). Adding the OTHER samples into the same space clearly indicated a continuous disease spectrum as well as a progression course along the trajectory (Figure 2B, D, and F), although the trajectory derived from HCN is less smooth. We evaluated how accurately the trajectory models AD’s progression, by linearly correlating the SI, i.e. the pseudotime each sample occupies along the trajectory from the starting point (presumably the normal state) to its final location, to its respective neuropathological and clinical biomarkers. We observed that SI correlated strongly with all AD specific biomarkers excluding diffuse plaques (P < 2.2E-16, R^2^ > 0.31, Figure 2G and H, Supplementary Table 4 and 5). Again, the regression metrics are the best for the model built from DLPFC followed by PCC, with HCN performs the least satisfactorily. All three models still considerably outperform our previous model built based a subset of the DLPFC samples[4].

**Figure 2.**
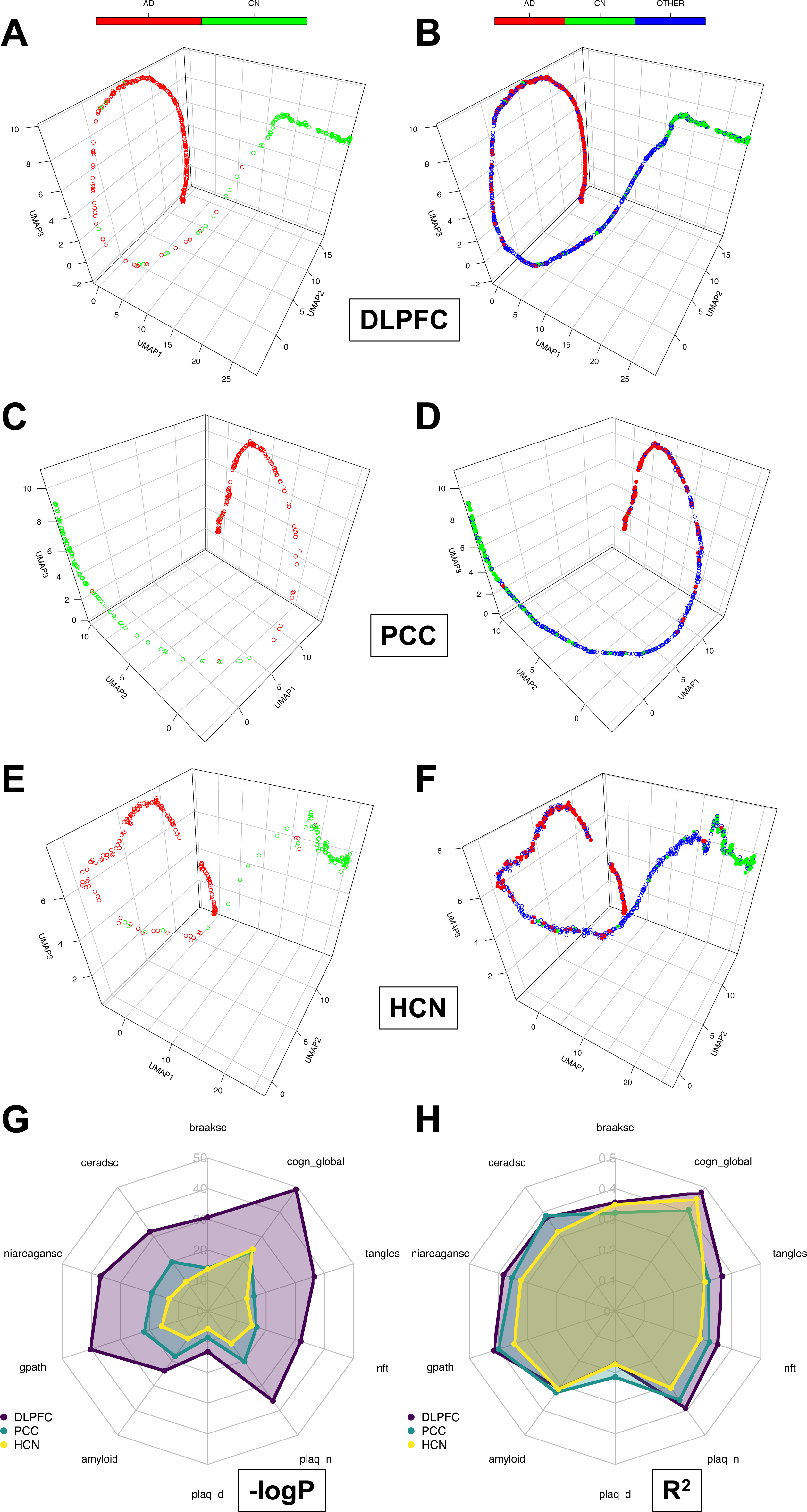
The pseudo-temporal trajectories from the trained models for the transcriptome from three regions of ROSMAP cohort and the SI correlation with phenotypical data. A-B): DLPFC without and with OTHER samples. C-D): PCC without and with OTHER samples. E-F): HCN without and with OTHER samples. G-H): Spider plots showing the linear correlations of SIs with phenotypical traits for all three models.

The three models were then applied individually to the harmonized transcriptomic data from both the MAYO and MSBB cohorts to derive their respective trajectories. Data from the MAYO cohort came from two different brain regions: TCX and CER. After projecting into the same 3D UMAP space, the subject distributions along the trajectories from the two different brain regions showed drastically different patterns (Figure 3A and B, DLPFC model only). For TCX, it showed the distributions of different locations for AD versus CN subjects along the trajectory similarly to those from ROSMAP data, while this was not observed for CER. This is confirmed by the results obtained from linear regression of the SI versus pathological biomarkers (Braak and Thal scores, Fig. 3C and D). Only in the TCX samples were the SIs found to be significantly correlated with both Braak (P = 7.98E-14, 4.80E-9 and 7.40E-6 respectively) and Thal scores (P = 2.95E-7, 1.04E-5, and 5.13E-3 respectively). Again, when applied to TCX dataset, the models explained a large amount of variance overall for both biomarkers, with correlation coefficient R in the range of 0.61 to 0.72 for the three models respectively. We also observed that the model from DLPFC is the most, while HCN is the least predictive of biomarkers in the MAYO cohort (Figure 3E and F, Supplementary Table 6). Interestingly, the models derived from DLPFC and PCC also showed some predictive power for Braak score in CER dataset (P = 4.94E-4 and 9.84E-3) although the PVEs are low (< 0.09 in comparison with > 0.29 in TCX data), indicating that tauopathy in AD is associated with widespread transcriptomic changes, including in regions comparatively spared from development of neurofibrillary tangles.

**Figure 3.**
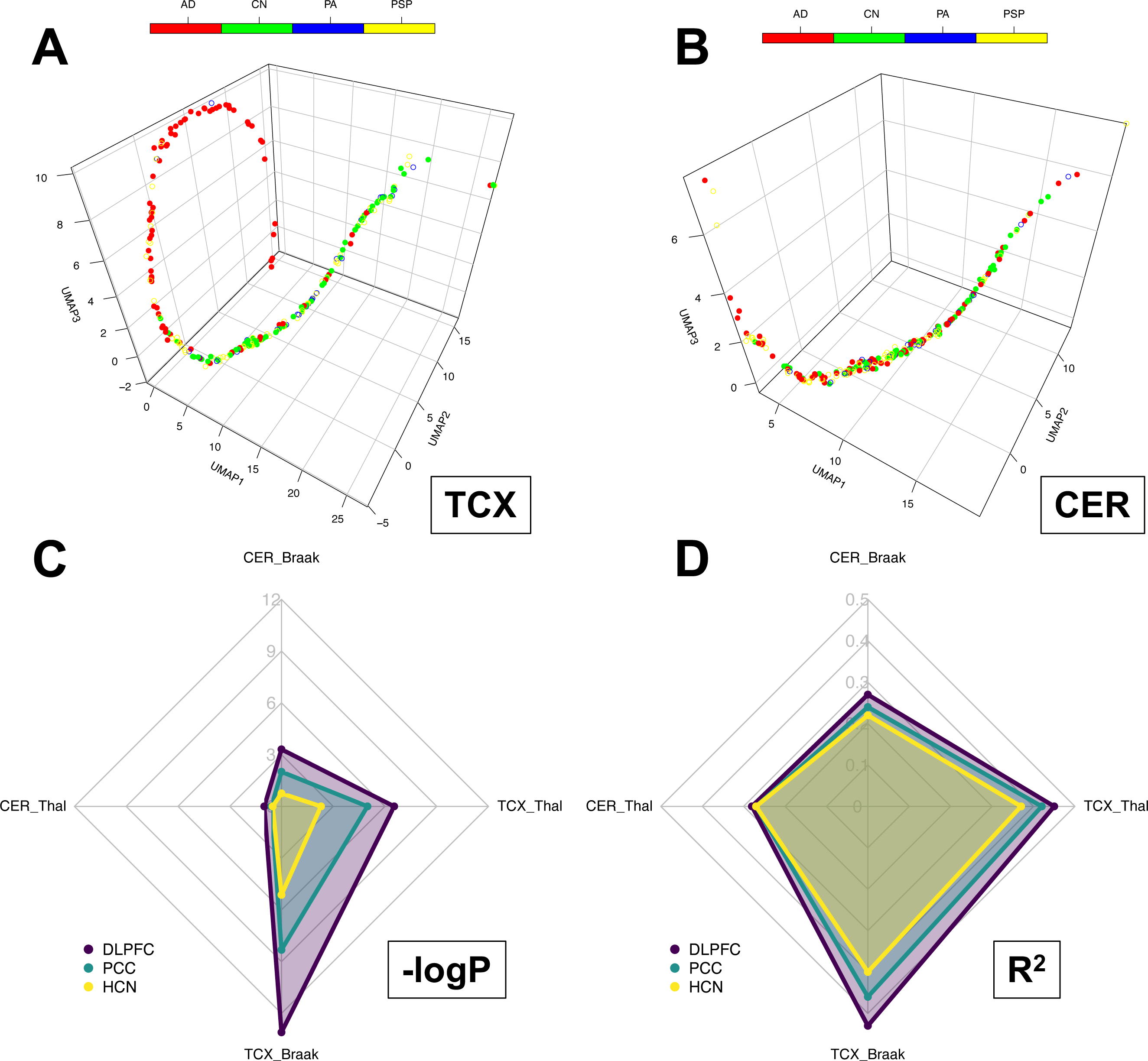
The pseudo-temporal trajectories from the forward pass of ROSMAP model and mapping to the same 3D space as ROSMAP (DLPFC only), based on the transcriptome from two regions of MAYO cohort and the SI correlation with phenotypical data from all three models’ predictions. PA: pathological aging; PSP: progressive supranuclear palsy. A) TCX mapped to DLPFC; B) CER mapped to DLPFC. C-D): Spider plots showing the linear correlations of SIs with phenotypical traits from all three models.

For the MSBB cohort, the models were applied to the gene expression profile of all four sampled regions [FP (BM10), STG (BM22), PHG (BM36), and IFG (BM44)] and all regions show similar albeit slightly different trajectories (Figure 4A-D, DLPFC model only), with the SI consistently significantly correlated with all the neuropathological and clinical biomarkers (Braak score, PlaqueMean, CDR scale and CERAD score, Supplementary Table 7), for the models from DLPFC and PCC. Still, the model from HCN doesn’t predict some of the biomarkers well when applied to certain brain regions such as Braak or CERAD score in FP (Figure 4E and F, Supplementary Table 7).

**Figure 4.**
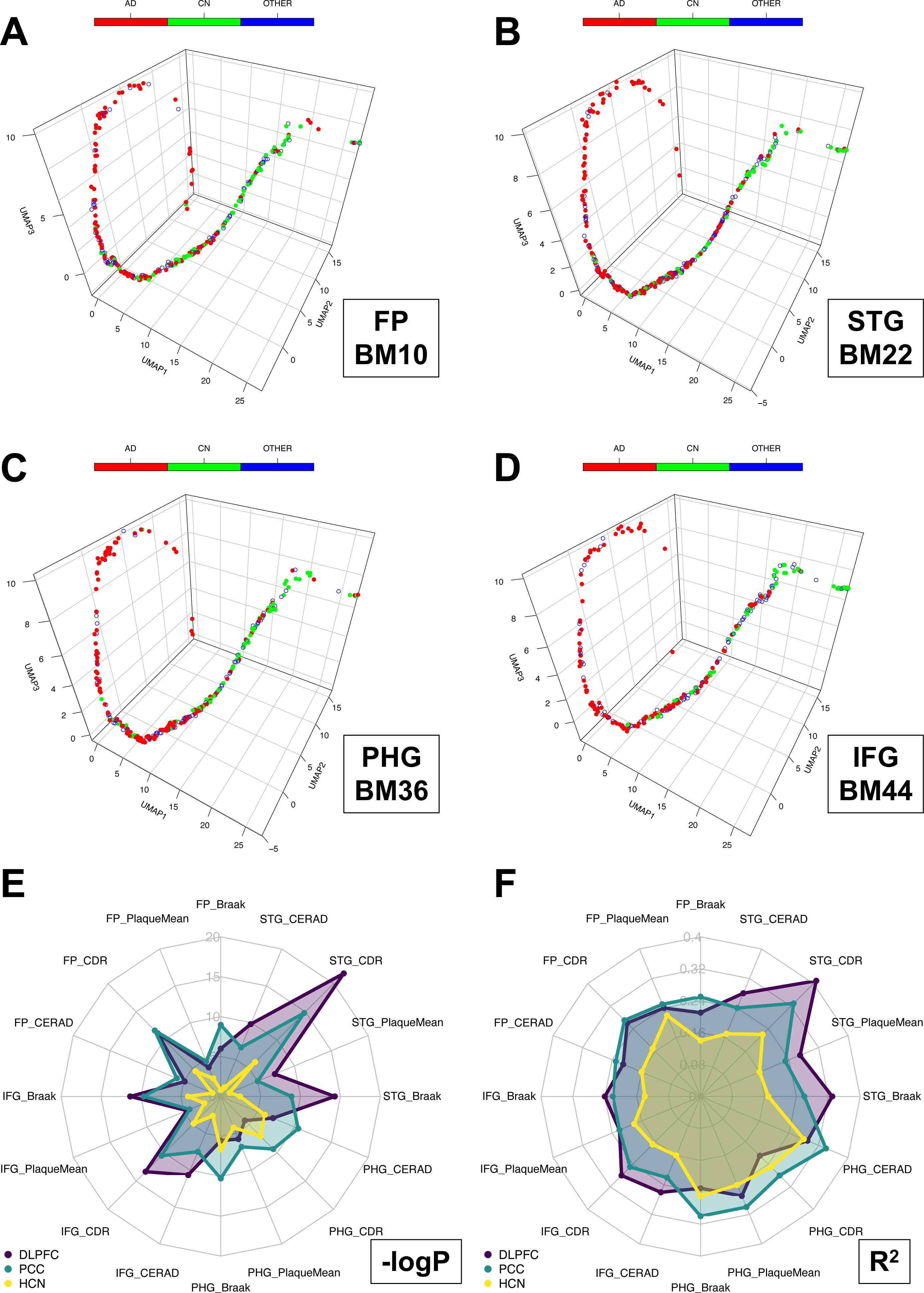
The pseudo-temporal trajectories from the forward pass of ROSMAP model and mapping to the same 3D space as ROSMAP (DLPFC only), based on the transcriptome from four regions of MSBB cohort and the SI correlation with phenotypical data from all three models’ predictions. A-D) Four regions (FP, STG, PHG, and IFG) mapped to DLPFC. E-F): Spider plots showing the linear correlations of SIs with phenotypical traits from all three models.

### 3.3 Comparison of the models trained from and applied to different brain regions

As reported in 3.1 and 3.2, when comparing the models trained from three different brain regions in ROSMAP, it is evident that neuropathology and cognitive impairment from AD are better reflected in the transcriptomes of cortical regions. The model from HCN shows relatively poor performance in classification metrics, aligning with phenotypical traits within the cohort, and prediction in external datasets. In contrast, models from DLPFC and PCC show comparably excellent performances, especially considering that PCC was trained with a smaller number of samples. When comparing the SIs derived from the three regions for the same subjects, we also observed that DLPFC-PCC pairs are most significantly correlated with each other (n = 621, p = 1.07E-142, R^2^ = 0.65, no other covariates considered), followed by DLPFC-HCN (n = 670, p = 8.27E-118, R^2^ = 0.55), and PCC-HCN (n = 466, p = 1.46E-90, R^2^ = 0.58). We subsequently focused the comparison of the models’ predictive performances from the DLPFC and PCC regions. When applied to external datasets, we observed that the model from DLPFC demonstrates the greatest predictive power and correlation coefficient of traits with the transcriptome from the TCX region of MAYO cohort, and IFG and STG regions of MSBB cohort, all of which are located at the either prefrontal cortex (DLPFC and IFG) or temporal lobe (TCX and STG). Conversely, the model from PCC is more predictive of data from PHG of MSBB cohort (Figure 3 and 4), both of which are deemed as part of the hippocampocentric subdivision of the paralimbic zone[35]. This illustrates that nuanced molecular changes in different brain regions affected by AD could be captured by sophisticated ML methods.

### 3.4 Index genes derived from the model interpretation

We set out to interpret our models using one of the latest model interpreting methods, the SHAP values, to reveal which features (ie genes) have the most impact (whether positively or negatively) on a specific prediction. We averaged out the feature contributions to AD and CN respectively, then took the absolute mean of the two to evaluate their importance to the classification. To robustly capture highly relevant genes, the training procedure was repeated 100 times using the same hyperparameters but different random number seeds, with the goal of simulating a “consensus network” and sampling as much space as possible. The process of averaging the SHAP values of all the features from each training to obtain the final SHAP metric and extract the list of the most salient genes is illustrated in Supplementary Figure 2.

Following this procedure, we selected 1,317, 1,594 and 1,643 genes for the models from three brain regions respectively (Supplementary Table 8A-C). They were labeled as “index genes” (IGs) for the subsequent in-depth analysis. We compared the IGs with those differentially expressed genes (DEGs) identified from the same datasets (syn26967457), by the same contrast of neuropathological confirmed AD vs CN comparison. There are some overlaps, but still considerable differences between the IGs and DEGs for the same brain region, as shown in Supplementary Figure 3A. Unlike those DEGs, IGs are not bounded by their fold changes or P values in the contrast, as in fact, the logFC and P values in the contrast for these IGs show a consistently even distribution (Supplementary Figure 3B and C), characterized by the coordinated regulations at molecular level.

We observed more overlaps between IGs of DLPFC and PCC as expected. All highly significant (P < 2.2E-16), the odds ratio (OR) of the overlaps between DLPFC and PCC (20.3) is higher than those between DLPFC and HCN (10.3) and PCC and HCN (15.5), confirming that the expression profiles of the two from cortical regions are more similar with each other than the subcortical region (HCN).

### 3.5 Co-expression modules of IG reveal molecular changes associated with AD from different brain regions

We derived co-expression network modules for the IGs based on the gene expression profiles of the whole cohort from the three regions respectively. For DLPFC, five distinct modules were resolved from the profile (Figure 5A-E), ranging from 78 to 542 genes in each module.

**Figure 5.**
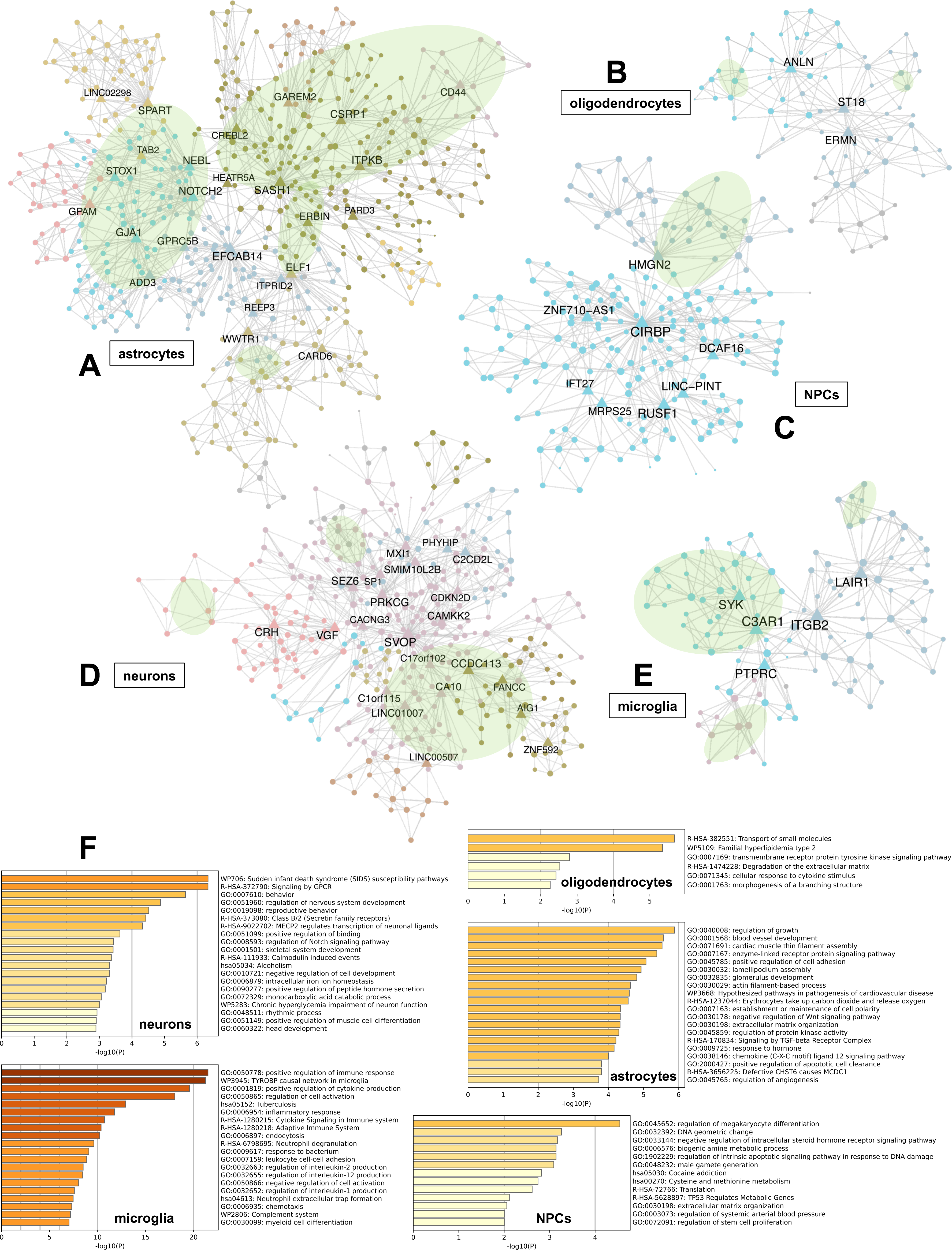
Co-expression modules resolved from the expression profiles of IGs in DLPFC region and their functional annotations. A-E) Five modules clustered by their cell type enrichment. Only hub genes are labeled. F) Functional enrichment for each module.

Functional annotations implicated them in different cellular processes from five major cell types (Figure 5F, Supplementary Table 8A). Most of the genes in the modules from microglia and oligodendrocytes are upregulated, while those from neurons are downregulated (Supplementary Figure 3B and C), indicating a consistent pattern of gliosis and neurodegeneration[36]. The wide range of gene regulation in astrocyte module also illustrates the cell type specific heterogeneity from their diverse functions in AD[37]. In contrast, we obtained 14 and 18 modules ranging from 25 to 227 genes for PCC and HCN profiles respectively (Supplementary Table 8B and C), some of which are subsets of the five modules from DLPFC. The more fragmented module structure could be attributed to the facts that the RNA qualities for PCC are significantly lower than DLPFC (mean RIN = 5.64 vs 6.26, P = 4.82E-18, Mann-Whitney U test), while the model from HCN doesn’t perform well. In addition, we have smaller numbers of the samples to train the two models than DLPFC. Consequently, there are very few common networks shared among the modules from all three regions. Nevertheless, we still observe sizeable submodules in common between the networks of DLPFC and PCC, especially for the networks from neurons, astrocytes, and microglia (Figure 5A, D and E), with at least two out of total three submodules in microglia, two submodules in astrocytes and two submodules neurons overlapped respectively (Supplementary Table 9).

We subsequently focused our analysis on the modules from DLPFC, for a comprehensive interpretation of the best model in our study to obtain the biological insights of molecular changes in one of the most important brain regions implicated in AD. We observed that the gene nodes are enriched for genetic risk loci[38, 39] from genome-wide association studies (GWAS) in the two glial modules (e.g. TREM2, MYO1E, PLCG2, CD33, HLA genes among others in microglia, P = 7.45E-6, OR = 8.75; and CR1, ADAMTS1, C2, C4A/B, IQCK among others in astrocytes, P = 2.87E-3, OR = 2.65, Supplementary Table 8A).

Interestingly, the microglia module depicts an extensive picture of immune response in AD (Figure 6A). In addition to the enrichment of GWAS loci, there is a high enrichment of disease associated microglia (DAM) signature genes[40] such as SPP1, ITGAX, CLEC7A, TMEM119, and TREM2 (P = 5.09E-12, OR = 20.36), as well as the TYROBP causal network in microglia (WP3625)[41] (P = 5.28E-26, OR = 182.9) and the CD33/LAIR-1 inhibitory networks[42]. In the center of the networks, as well as from an overlapped module in PCC, is the hub gene C3AR1, recently identified as a major player mediating neuroinflammation and tau pathology in AD[43, 44]. It is notable that many of these genes (e.g. PLCG2, MYO1E, VSIG4, LAIR1 and HLA genes) are not identified as DEGs in the same datasets used in our study by falling short of logFC or P value cutoff, although they have been reported as differentially expressed in single-nucleus RNA sequencing (snRNA-seq) data[45, 46], demonstrating the power of deep learning methods to uncover nuanced signals in convoluted data.

**Figure 6.**
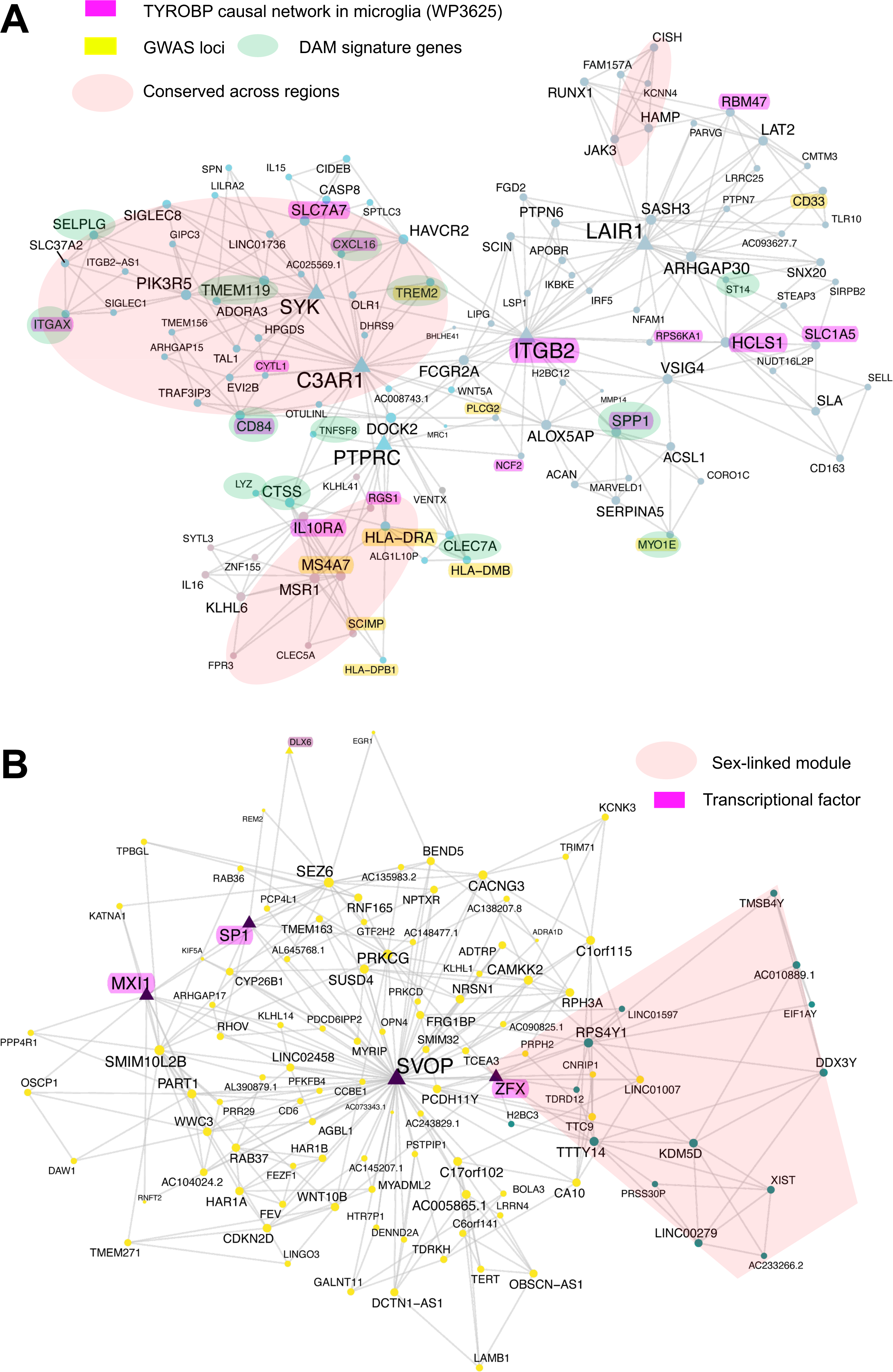
Curated co-expression plots for two modules from DLPFC region with all gene nodes labeled. A) Microglia module. B) A subset from the neuron module including the hub gene SVOP and a sex-linked submodule.

The astrocyte module presented the largest overlaps between DLPFC and PCC regions from our models (Figure 5A). Among them, we identified the signatures from the GJA1 centered genetic networks[47] (P = 3.41E-27, OR = 7.62), the disease associated astrocytes (DAA) such as GFAP, LGMN, C4A/B [48] (P = 9.46E-7, OR = 5.86) and other astrocyte signatures obtained from snRNA-seq data [49]. Some hubs of the networks are also novel AD risk gene (SASH1) identified using machine learning GWAS platform[50] or known risk gene (ITPKB) for other neurodegenerative diseases such as Parkinson’s disease (PD)[51] but also implicated in AD[52].

### 3.6 Sex-linked module and transcriptional factor (TF) in DLPFC neurons

We observed that the neuron module possesses several key gene hubs such as CACNG3, VGF, NPTX2, RPH3A, SVOP, and BDNF, recently reported to exhibit positive associations with global cognitive function and negative associations with neuropathology across various excitatory neuron subtypes in AD (Figure 5D). They were found to be within a consensus signature significantly associated with global cognitive function in at least three different excitatory neuron subtypes, and prominently linked to both pre- and postsynaptic compartments, from a comprehensive analysis of snRNA-seq data from the prefrontal cortex (PFC) brain region within the ROSMAP cohort[53]. Again, they are not completely recapitulated by DEG analysis from bulk tissue RNA-seq data.

Our data show one prominent hub gene, which interconnects many of the aforementioned hubs, synaptic vesicle 2 related protein (SVOP), a protein involved in synaptic vesicle transport.

Intriguingly, SVOP is also connected with a co-expression submodule primarily composed of sex-linked genes such as XIST, TTTY14, and KDM5D (Figure 6B), by the X-linked zinc finger transcriptional regulator ZFX, which is known to escape X chromosome inactivation (XCI)[54, 55]. We assessed the transcriptional regulation of SVOP using the expression profile of DLPFC neuron module, and identified the candidate TFs that regulates SVOP expression (Supplementary Table 10). Among them, three TFs (SP1, MXI1, and ZFX) are directly connected to SVOP in the module (Figure 6B), all of which would be expected to act as transcriptional repressors based on their inverse expression correlation to SVOP.

Since ZFX and ZFY are X-Y homolog pairs (gametologs) assumed to have the same function, we examined the contribution of the two sex-linked genes to the expression of SVOP, for the whole cohort from DLPFC region. A simple linear regression shows that the expression of both genes is significantly correlated with SVOP expression, although ZFY at a much lesser degree (Figure 7A, Supplementary Table 11). In addition, both correlations show considerable residual sex effects in the model. To account for their additive effects, we reprocessed the dataset with raw gene counts (syn22231797), adding an additional feature by combining ZFX+ZFY gene counts. Indeed, we observed that the cumulative expression of ZFX and ZFY completely explains the sex differences of SVOP expression by inverse correlation, with no residual sex effect, or additional diagnosis difference unaccounted for in the linear model (Figure 7A, Supplementary Table 10). Nevertheless, the cumulative TF expression is still lower in male, due to the escape of XCI of ZFX in females and lower expression of ZFY in males in comparison with their homolog ZFX expression[56]. Consequently, SVOP is more significantly downregulated in AD females than their male counterparts (Figure 7B). As SVOP plays a central role in the co-expression module associated with neuronal loss in AD, our research provides direct evidence of the molecular mechanisms underlying sex chromosome involvement and sexual dimorphism in AD.

**Figure 7.**
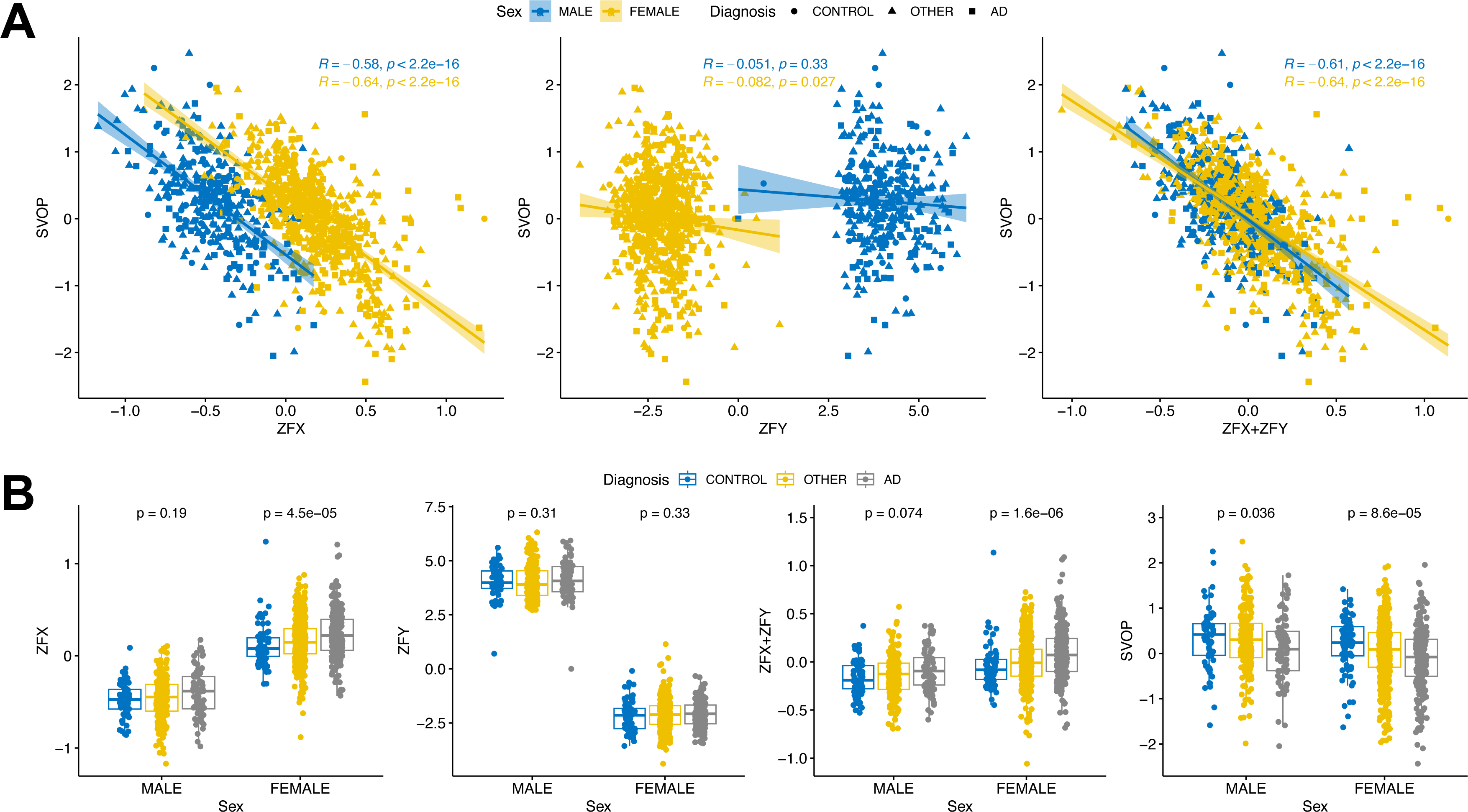
SVOP and ZFX (ZFY) co-expression based on the reprocessed ROSMAP data from DLPFC region. A) Regression between SVOP and ZFX, ZFY and their combined expression. B) Boxplots showing ZFX, ZFY, their cumulation and SVOP expression stratified by sex.

## 4. Discussion

In this study we present a comprehensive interpretable deep learning framework on the RNA-seq data obtained from multiple postmortem brain regions of the ROSMAP cohort, a large clinical cohort of AD. We also applied the trained models to the transcriptomic data from two independent cohorts, the MAYO and MSBB cohort. Our models show excellent predictive power in aligning the transcriptomes with clinical and neuropathological traits in both internal and external validations, as indicated by the model metrics from the SI for delineating the progressive pseudotemporal trajectories in each dataset. This underscores the broader applicability of the framework in the study of neurodegenerative diseases such as AD as a continuum[57].

By individually modeling the transcriptomes from three distinct brain regions of the ROSMAP cohort, we demonstrate the capacity of deep learning methods to learn and capture subtle distinctions present in diverse brain regions affected by AD. The models successfully differentiate the non-specific subcortical region in AD, i.e. HCN. Moreover, they distinguish nuanced variations in the data from the two cortical regions (DLPFC vs. PCC), evident in the observable differences in predictive accuracy when applied to data from other relevant regions in an external cohort (MSBB, Figure 4, Supplementary Table 7). This reiterates the notion that molecular changes in AD-affected brains are both specific and regional, underscoring the importance of considering these factors in comprehensive studies[58].

The most pivotal insights derived from the study come from the IGs through model interpretation. By applying the SHAP method based on cooperative game theory to explain the outcome of the model, our approach provides a way to fairly allocate contributions of each feature, with both global and local interpretability. The summarized importance score would therefore offer an overview of feature importance across the entire dataset for AD vs control classification. Our framework consistently excels in extracting input features that go beyond the constraints of logFC or P values stemmed from traditional DEG analysis, thus revealing nonlinear, coordinated, and cell type specific gene regulation from bulk tissue data. These signals are otherwise only available from deconvoluted or higher resolution omics data, such as single cell RNA-seq. It would eventually be desirable to apply the framework to such kind of data to obtain even more novel biological insights for disease etiology at cell level.

The co-expression networks derived from the IGs from the two cortical regions, particularly DLPFC highlight the critical roles of gliosis and neurodegeneration in AD. In microglia, the networks corroborate many of the established activated pathways implicated in AD, such as TYROBP causal network and the DAM signatures. In particular, the networks draw attention to the significance of the TREM2-DAP12-SYK pathway, which coordinates neuroprotective microglial responses in AD[59, 60]. It will be worthwhile to further investigate the relationship of this pathway and the C3 and C3A receptor (C3AR1) signaling given their close co-expression pattern. The data highlights the neuroimmune axis as evidenced by MHC class II signaling transduction, which suggests an intricate interplay of adaptive and innate immune systems both within and outside the brain influencing the etiology and pathogenesis of AD[61]. It is worth noting that the networks also implicate the role of the C1Q/CD33/LAIR-1 inhibitory complex in AD, and it seems it is only present in DLPFC. Since the complex has been reported to dampen monocyte immune response[62], in-depth study is thus warranted to explore the functions of this complex and its influence the balance between immune activation and tolerance in AD, and whether this effect is brain region specific.

It has now been widely acknowledged that AD disproportionately affects women in both disease prevalence and rate of symptom progression, but the mechanisms underlying this sexual divergence are still being actively pursued. From transcriptional analyses, gene dysregulation in AD is particularly prominent in the neuronal cell populations, especially in the females[63].

Notably, for almost all the hub genes identified in the neuronal module in this study such as CACNG3, VGF, NPTX2, RPH3A, SVOP, and CA10, their fold changes in DE analysis are more significant in the females than males by sex stratified analysis (syn26967458), which suggests a more severe neuronal damage in females. Conversely, the transcriptional factors predicted to repress the expression of SVOP (SP1, MXI1 and ZFX) all show more pronounced upregulation in females in DLPFC (Supplementary Figure 4). Recently it has been reported that the human Y and inactive X chromosomes similarly modulate autosomal gene expression, with the homologous transcription factors – ZFX and ZFY acting in a mutually and cumulatively dose-dependent fashion[64]. Most importantly, they are prioritized as one of the genes on sex chromosomes most likely to contribute to male-female differences in common disease[65]. We observed highly correlated co-expression between their cumulative expression and that of SVOP, with the sum greater in females and more significantly upregulated in AD. Together with the two other transcriptional factors (SP1 and MXI1), their expressions explain over 70% variances of the expression of SVOP in the DLPFC data, indicating a highly probable orchestrated transcriptional regulation.

SVOP is a member of the synaptic vesicle glycoprotein 2 (SV2) family primarily associated with synaptic vesicles. It plays a role in the regulation of neurotransmitter release at synapses[66] although there is limited study for its characterization. Its expression has been found to be positively correlated with cognitive function and consistently downregulated in multiple neuron subtypes from AD brains especially in females, in numerous transcriptomic profiles by snRNA-seq[53, 67–69]. Further characterization of the transcript and protein and their roles in neurological function and disorder is thus warranted.

Although still significant, the co-expressions of SVOP with the TFs in the two other brain regions are not as strong, with residual sex effect not completely modeled (Supplementary Table 10). It remains to be further investigated the details of the epigenomic landscape of the TFs around SVOP in different cell types from different AD affected brain regions, and how and why their regulations are disrupted in AD differently in the two sexes. In this work we report a novel mechanism that illuminates how sex chromosomes impact upon AD transcriptomics in a sex-specific manner. It therefore provides valuable preliminary data for further research on the biological mechanism(s) of action and the implications for disease pathogenesis of sex differences in AD.

Among the gene nodes in the sex-linked submodule, excluding the female sex marker XIST, all the other sex-linked genes (RPS4Y1, LINC00279, DDX3Y, TMSB4Y, TTTY14, AC010889.1, KDM5D, and EIF1AY) are on the Y chromosome. Their joint co-expression presence is most likely due to the distribution of sex effect to several highly expressed Y-linked genes, since they are only linked to the parent module by ZFX. Yet there is another Y-linked gene PCDH11Y showing co-expression with several genes in the neuronal module including ZFX, SLC22A25, and SVOP (Figure 6B). It is notable that its X-homolog PCDH11X has been reported to be associated with higher risks of developing AD in women, although the finding cannot be replicated in subsequent studies[70], leaving the window open for more research. In the future we would also anticipate reanalyzing this data accounting for the differential presence of XIST and Y-linked gene expression in genotypic male and female samples by sex chromosome complement (SCC) aware alignment approach such as XY_RNAseq[71]. Given that ZFX in females and ZFX+ZFY in males gives a robust signal across all samples, our work highlights the importance of considering sex chromosome shared genes (gametologs) like ZFX and ZFY, which have conserved function uniquely when incorporating the sex chromosomes[64].

One limitation of our study is that we applied the framework on the transcriptomes obtained from bulk-tissue which could be affected by cell type proportion during disease progression. In light of the ever-growing corpus of single cell multi-omics data and the demonstrated capability for our approach to disentangle highly convoluted data, we anticipate these methods will offer broad utility in advancing our understanding of complex human diseases such as AD.

## Figure Legend

Supplementary Figure 1. Receiver operating curves (ROCs) for the three trained models.

Supplementary Figure 2. The process of training 100 models and obtaining the final SHAP values for feature selection.

Supplementary Figure 3. The comparisons of the IGs and DEGs of ROSMAP. A) An upset plot comparing the IGs and DEGs for all three regions. B) The IGs distribution on the Volcano plot from DLPFC DEG analysis (data from syn26967457). Each module is colored respectively. C) Boxplots for the IGs distribution by their -logP and log_2_FC in the DEG analysis. Color scheme is the same as B).

Supplementary Figure 4. Heatmap showing the log_2_FC for the IGs in the neuronal module (377 genes) in AD vs control comparison by DEG analysis (data from syn26967458) stratified by sex. Data is sorted by the middle panel from joint analysis. The sex-linked genes are labeled on the left. Other important genes are labeled on the right. DEG cutoffs for the joint analysis (|log_2_FC| > 0.263) are marked.

## Acknowledgements

The results published here are in whole or in part based on data obtained from the AD Knowledge Portal (https://adknowledgeportal.org). The RNA-seq Harmonization Study was supported by the NIA grants U01AG046152, U01AG046170, U01AG046139 and U24AG061340. Study data in ROSMAP cohort were provided by the Rush Alzheimer’s Disease Center, Rush University Medical Center, Chicago, IL, USA. Data collection was supported through funding by NIA grants P30AG10161 (ROS), R01AG15819 (ROSMAP; genomics and RNA-seq), R01AG17917 (MAP), R01AG36836 (RNA-seq), the Illinois Department of Public Health and the Translational Genomics Research Institute. Additional phenotypic data were requested at https://www.radc.rush.edu. The data for MSBB cohort were generated from post-mortem brain tissue collected through the Mount Sinai VA Medical Center Brain Bank and were provided by Dr Eric Schadt from Mount Sinai School of Medicine. The MSBB study was led by Dr Nilufer Ertekin-Taner and Dr Steven G. Younkin, Mayo Clinic, Jacksonville, FL, USA, using samples from the Mayo Clinic Study of Aging, the Mayo Clinic Alzheimer’s Disease Research Center and the Mayo Clinic Brain Bank. Data collection was supported through funding by NIA grants P50AG016574, R01AG032990, U01AG046139, R01AG018023, U01AG006576, U01AG006786, R01AG025711, R01AG017216, R01AG003949, NINDS grant R01NS080820, CurePSP Foundation and support from Mayo Foundation. Study data include samples collected through the Sun Health Research Institute Brain and Body Donation Program of Sun City, Arizona. The Brain and Body Donation Program is supported by the NINDS (U24NS072026 National Brain and Tissue Resource for Parkinson’s Disease and Related Disorders), the NIA (P30AG19610 Arizona Alzheimer’s Disease Core Center), the Arizona Department of Health Services (contract 211002, Arizona Alzheimer’s Research Center), the Arizona Biomedical Research Commission (contracts 4001, 0011, 05-901 and 1001 to the Arizona Parkinson’s Disease Consortium) and the Michael J. Fox Foundation for Parkinson’s Research. The authors acknowledge Research Computing at Arizona State University for providing computing resources that have contributed to the research results reported within this paper.

