## Supplementary figures and images for "Interpretable deep learning framework for understanding molecular changes in human brains with Alzheimer’s disease: implications for microglia activation and sex differences"

### Supplementary Figure 1

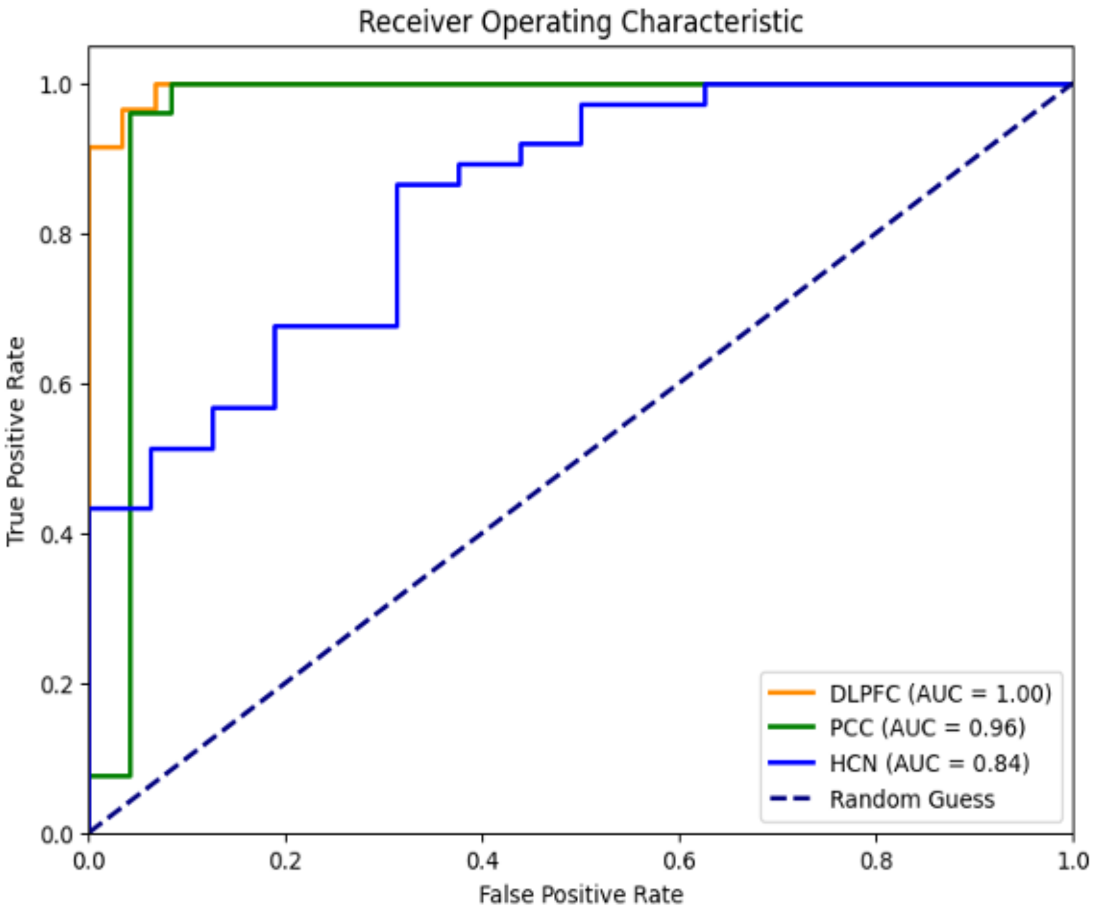

### Supplementary Figure 2

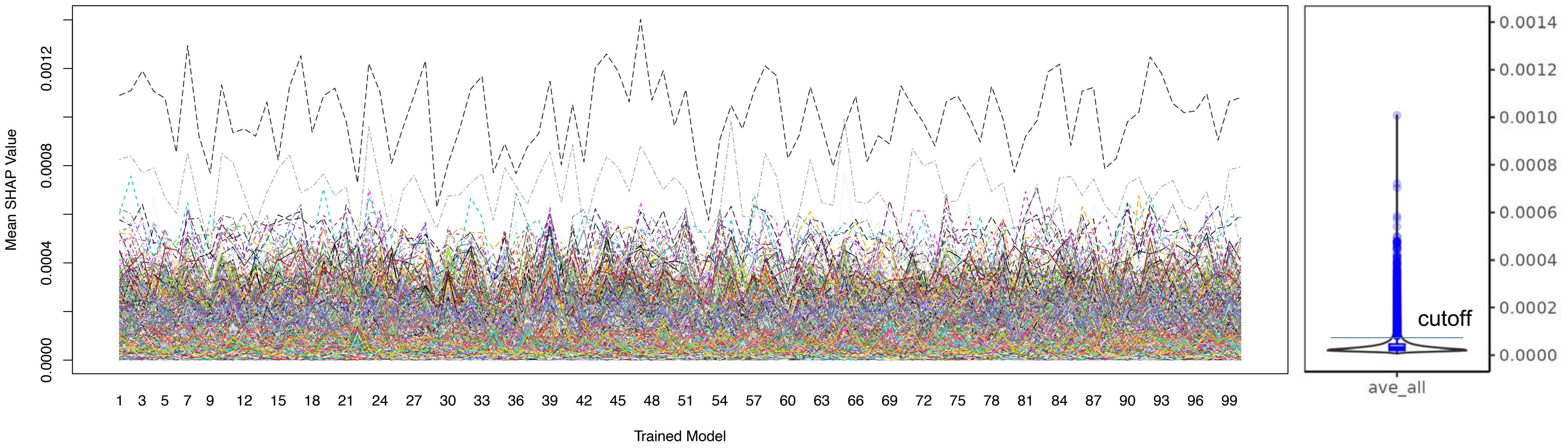

### Supplementary Figure 3

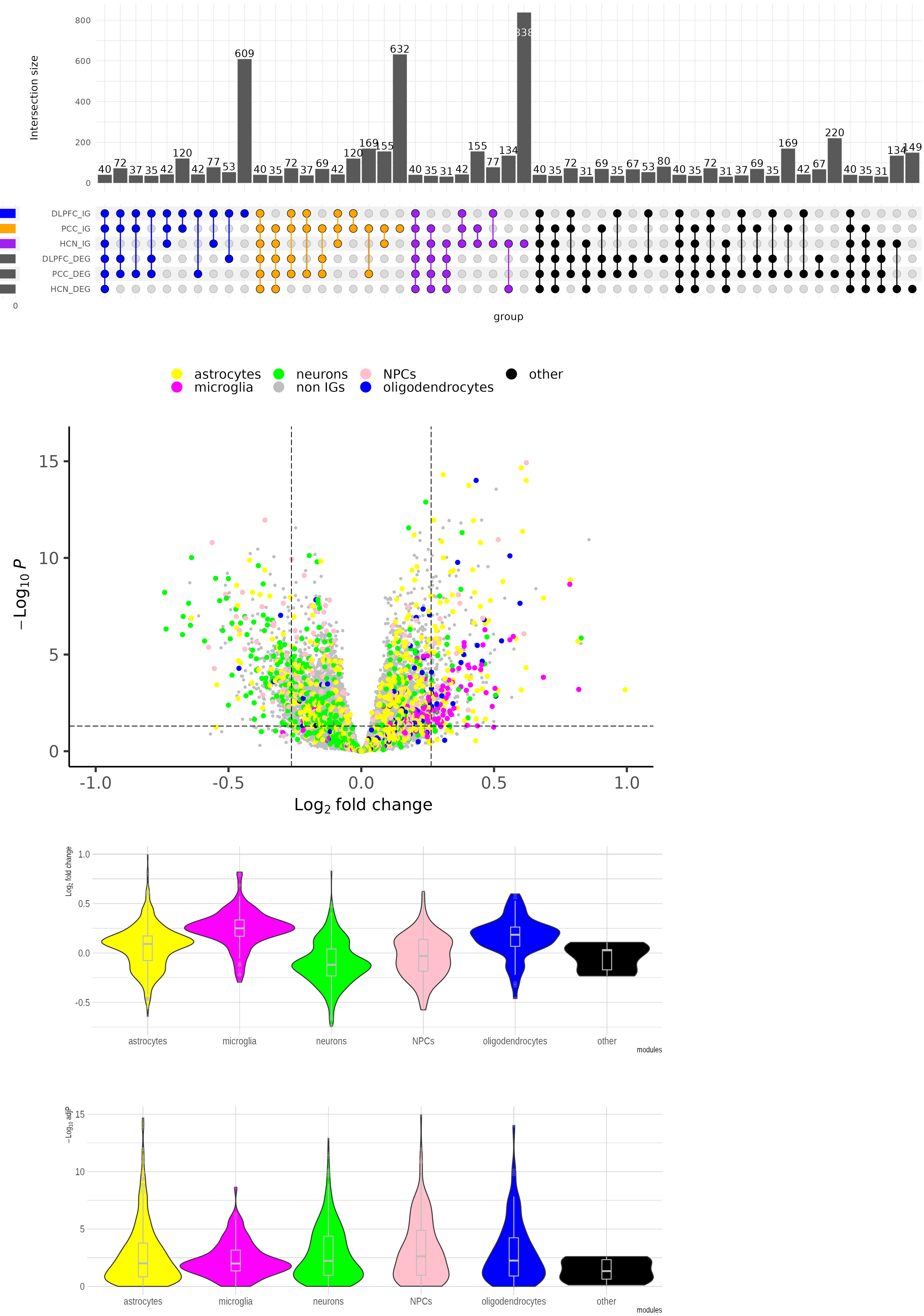

### Supplementary Figure 4

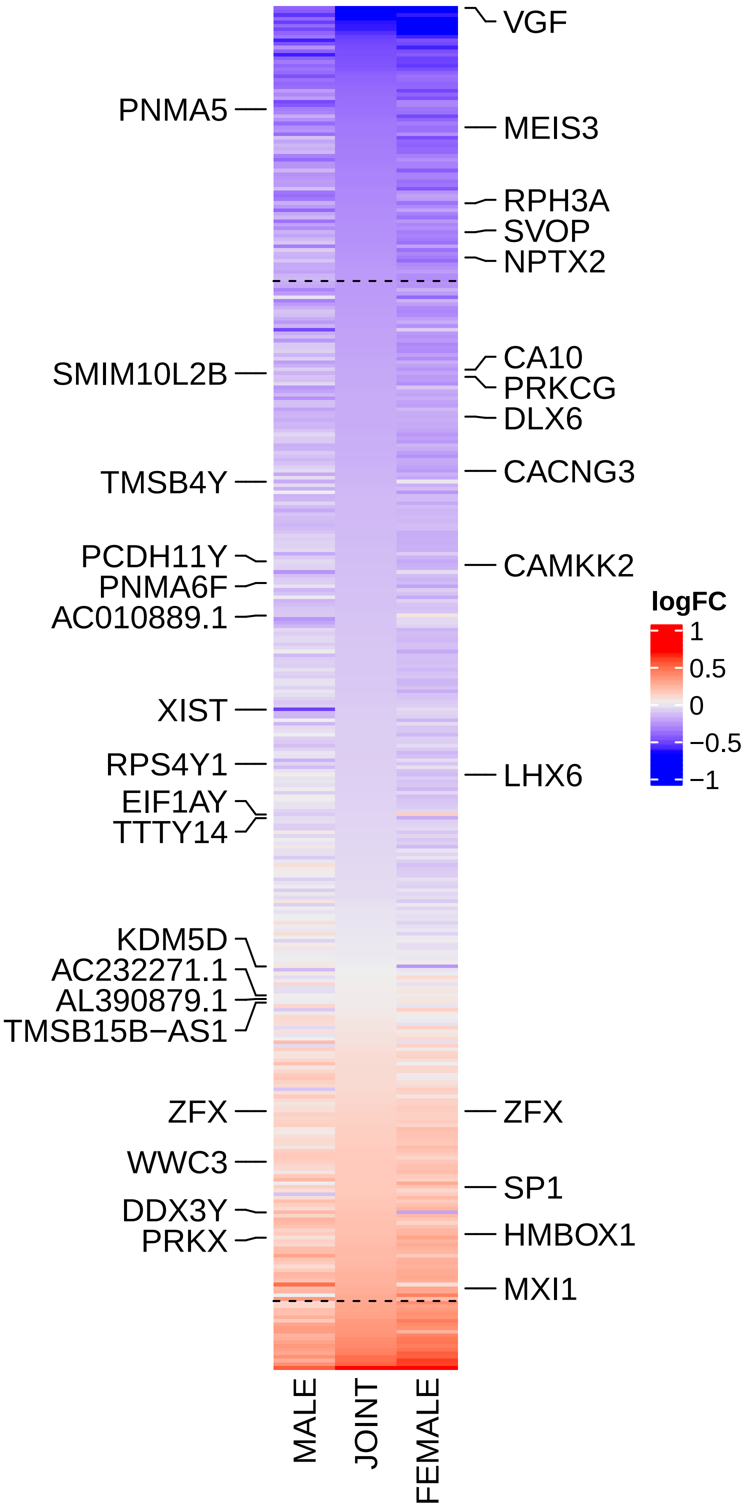
